# Combining Aperiodic 1/f Slopes and Brain Simulation: An EEG/MEG Proxy Marker of Excitation/Inhibition Imbalance in Alzheimer’s Disease

**DOI:** 10.1101/2022.12.21.521529

**Authors:** Pablo Martínez-Cañada, Eduardo Perez-Valero, Jesus Minguillon, Francisco Pelayo, Miguel A. López-Gordo, Christian Morillas

## Abstract

Accumulation and interaction of amyloid-beta (Aβ) and tau proteins during progression of Alzheimer’s disease (AD) are shown to tilt neuronal circuits away from balanced excitation/inhibition (E/I). Current available techniques for noninvasive interrogation of E/I in the intact human brain, e.g., magnetic resonance spectroscopy (MRS), are highly restrictive (i.e., limited spatial extent), have low temporal and spatial resolution and suffer from the limited ability to distinguish accurately between different neurotransmitters complicating its interpretation. As such, these methods alone offer an incomplete explanation of E/I. Recently, the aperiodic component of neural power spectrum, often referred to in the literature as the ‘ 1/f slope’, has been described as a promising and scalable biomarker that can track disruptions in E/I potentially underlying a spectrum of clinical conditions, such as autism, schizophrenia, or epilepsy, as well as developmental E/I changes as seen in aging. Using 1/f slopes from resting-state spectral data and computational modelling we developed a new method for inferring E/I alterations in AD. We tested our method on recent freely and publicly available electroencephalography (EEG) and magnetoencephalography (MEG) datasets of patients with AD or prodromal disease and demonstrated the method’s potential for uncovering regional patterns of abnormal excitatory and inhibitory parameters. Our results provide a general framework for investigating circuit-level disorders in AD and developing therapeutic interventions that aim to restore the balance between excitation and inhibition.

## 1. Introduction

Abnormal accumulation and circuit-level interactions of amyloid-beta (Aβ) and tau proteins in the cortex are the neuropathologic hallmarks of Alzheimer’s disease (AD), starting decades before the emergence of clinically measurable cognitive deficits (Busche and Hyman 2020; Harris et al. 2020; Palop and Mucke 2016). Previous findings suggest that neuronal and circuit level modes of dysfunction (i.e., hyper- and hypoexcitability) can arise from the neuronal effects mediated by AD-related proteins. Interestingly, Aβ and tau have shown to exhibit opposing effects on local circuit dynamics. There is strong evidence that Aβ plaques, as well as soluble forms of Aβ, are a key player in driving neuronal hyperexcitability in AD, which might ultimately give rise to epileptiform activity (Zott et al. 2019; Busche et al. 2015; Verret et al. 2012; Busche et al. 2008). In direct contrast to Aβ-mediated effects, tau is associated with suppression of neuronal activity (hypoexcitability) and progressive loss of connectivity (Busche et al. 2019; Green et al. 2019). Moreover, Aβ and tau may not act independently but recent evidence suggests that both pathologies have synergistic effects (Busche and Hyman 2020). During AD progression, Aβ and tau show different temporal evolution profiles (in which tau pathology is delayed), are initially deposited in different brain regions (Aβ plaques are particularly found in medial prefrontal and medial parietal regions, while tau aggregates, in the medial temporal lobe) and propagate across other cortical regions as the disease progresses (Harris et al. 2020; Busche and Hyman 2020; van der Kant, Goldstein, and Ossenkoppele 2020; Leuzy et al. 2019; Palop and Mucke 2016). Given the complexity of the competing and synergistic functional effects between AD-related proteins, disambiguating their influence on neural dynamics and identifying how they modulate circuit excitation and inhibition (E/I balance) remains a topic of intense research (Ranasinghe et al. 2022; Maestú et al. 2021; Harris et al. 2020).

Direct and indirect in-vivo measurement of E/I is especially challenging in the human brain. Current noninvasive assessments of E/I in humans are largely restricted to magnetic resonance spectroscopy (MRS) measurements of in vivo concentrations of primary excitatory (glutamate) and inhibitory (γ-aminobutyric acid, GABA) neurotransmitters (Cuypers and Marsman 2021; Rideaux 2021; Steel et al. 2020; Stanley and Raz 2018). This approach presents several methodological challenges: the relatively low signals of glutamate and GABA complicate their interpretation because of their overlap with signals from other metabolites, it has low temporal and spatial resolution, and their measurements are usually restricted to one or few brain regions. For these reasons, current MRS approaches have limited utility for tracking variations of E/I across cortical regions or behavioural states. Recently, a range of novel features derived from electroencephalography (EEG) and magnetoencephalography (MEG) recordings have been described as robust proxy makers for noninvasive real-time measurements of changes in E/I balance (Ahmad et al. 2022; Manyukhina et al. 2022; Pablo Martínez-Cañada, Noei, and Panzeri 2021; Molina et al. 2020; Gao, Peterson, and Voytek 2017). One of the most promising candidate E/I biomarkers is the exponent of the 1/f spectral power law, often referred to in the literature as the ‘1/f slope’. Converging evidence from neuroimaging, pharmacological, chemogenetic and computational modelling studies has linked changes in this marker to conditions related to altered E/I balance (Pani, Saba, and Fraschini 2022), such as autism (Manyukhina et al. 2022; Trakoshis et al. 2020), schizophrenia (Molina et al. 2020), epilepsy (van Heumen et al. 2021; Pani et al. 2021) and attention-deficit/hyperactivity disorder (Karalunas et al. 2022; Robertson et al. 2019). The 1/f slope has been additionally shown to exhibit strong changes both across healthy aging (Tran et al. 2020; Voytek et al. 2015) and during early development (Chini, Pfeffer, and Hanganu-Opatz 2022; Schaworonkow and Voytek 2021), which have been associated to alterations in the relative E/I ratio.

Here, we introduce an approach that combines 1/f slopes from resting-state spectral data and simulation of a cortical microcircuit model with recurrent interactions between excitatory and inhibitory neuronal populations (P. Martínez-Cañada et al. 2021; Mazzoni et al. 2015; Mazzoni et al. 2008) for inferring E/I alterations in AD. We used our model-based inference approach to interrogate two recent publicly available datasets of EEG (Meghdadi et al. 2021) and MEG (Vaghari et al. 2022) recordings of patients with AD or Mild Cognitive Impairment (MCI). Finally, we assessed the potential of our approach for revealing the specific shifts in E/I occurring across cortical regions distinctively associated with tau and Aβ depositions at different stages of AD.

## 2. Materials and Methods

### 2.1. Participants

We used resting-state EEG and MEG data of people with AD or prodromal disease from open and publicly available datasets. The EEG dataset (Meghdadi et al. 2021) included recordings of individuals between the ages of 40-90 that were collected at four sites in the United Sates: Advanced Brain Monitoring (ABM) in Carlsbad, California; Advanced Neurobehavioral Health (ANH) in San Diego, California; Massachusetts General Hospital (MGH) in Boston, Massachusetts; and Mayo clinic (MAYO) in Rochester, Minnesota. Healthy controls were in two age groups: between 40-60 (HC2, n = 62) and between 60-90 (HC3, n = 52). Participants in the AD group (n = 26) were in the age range of 58-90. Patients were diagnosed with AD according to neurological evaluation and neuropsychological testing based on criteria developed by the National Institute of Neurological and Communicative Disorders and Stroke and the Alzheimer’s Disease and Related Disorders Association (NINCDS-ADRDA) (McKhann et al. 1984) and based on the Diagnostic and Statistical Manual of Mental Disorders (DSM-5) criteria for Major Neurocognitive Disorder (i.e., dementia) due to Alzheimer’s disease. Participants were excluded if they reported known neurological or psychiatric disorders, cardiac arrhythmias, heart failure (e.g., myocardial infarction), epilepsy, HIV+ diagnosis, bipolar disorder, or major depression (see ref. (Meghdadi et al. 2021) for more details).

MEG data was acquired from the BioFIND project (Vaghari et al. 2022), which included individuals with clinically diagnosed MCI and healthy controls pooled over a number of different projects from two sites: the MRC Cognition & Brain Sciences Unit (CBU) in Cambridge, England, and the Centre for Biomedical Technology (CTB) in Madrid, Spain. The MCI diagnosis of patients from CBU included Addenbrooke’s Cognitive Examination (ACE) and ACE Revised (ACE-R), and Mini Mental State Examination (MMSE) tests as standard. Positron Emission Tomography (PET) and fluidic biomarkers were not performed to patients, although Magnetic Resonance Imaging (MRI) was used for clinical follow-up in support of the diagnosis. The MCI diagnosis of patients from CTB was determined with intermediate probability according to the National Institute on Aging-Alzheimer Association criteria (Albert et al. 2011), i.e., given by a clinician based on clinical and cognitive tests, self- and informant-report, and in the absence of full dementia or obvious other causes. For some patients, there was additional biomarker evidence of atrophy from MRI or long-term follow up and genotyping for the APOE ε4 allele. For a subset of MCI patients, additional information was provided indicating whether or not they subsequently progressed to dementia (‘probable AD’) over subsequent years, according to their managing clinician. We only used MCI patients from this subset so that our results can be interpreted in terms of neural changes associated with probable AD (converted) or with patients who later did not convert to a dementia diagnosis (not converted). Additionally, and provided that age is the most important risk factor for developing Alzheimer’s Disease (Maestú et al. 2021; Harris et al. 2020), we limited our study of the BioFIND dataset to elder individuals (older than 75 years), including n = 50 MCI patients (n = 27 converted, n = 23 not converted) and n = 51 controls.

### 2.2. M/EEG preprocessing and power spectral analysis

We used the same preprocessing methods as described in the respective publications of the MEG and EEG datasets (Meghdadi et al. 2021; Vaghari et al. 2022). Briefly, EEG and MEG recordings were gathered during resting state with eyes closed and bandpass filtered (1-49 Hz for EEG data and from 0.5 to 48 Hz for MEG data). Bad channels of EEG data were detected and removed, as well as sources other than brain (e.g., eye blinks). EEG data were then epoched into one-second non-overlapping windows at each channel. The MEG analysis pipeline included MEG maxfiltering, epoching (two-second window), bad epoch detection, beamforming and region-of-interest (ROI) extraction (further details can found in ref. (Vaghari et al. 2022)).

EEG Power spectral densities (PSD) were available out of the box in the EEG dataset (Meghdadi et al. 2021). EEG PSDs were computed using LabX EEG processing software (Advanced Brain Monitoring Inc., Carlsbad, California). LabX employs modified periodogram PSD estimates with a 1-second-long Kaiser window (b = 6) and 50% overlap (see ref. (Meghdadi et al. 2021) for further details). We computed MEG PSDs of ROIs by ourselves using a similar approach. MEG PSDs were defined by time averaging spectrograms computed with a 1-second-long Kaiser window (b = 6) and 50% overlap. We normalised PSDs by dividing each EEG channel’s or ROI’s PSD by its mean power.

We parameterized power spectra into periodic and aperiodic components using the FOOOF algorithm (Donoghue et al. 2020) with the following configuration: frequency range = (1, 40) Hz, maximum number of peaks = 3, peak threshold = 1, peak width limits = (2, 50) Hz. In this study we disregarded the periodic (oscillatory) properties of spectral fittings. Aperiodic activity is described by a 1/*f^χ^* distribution, with exponentially decreasing power across increasing frequencies. When measured in the log-log space, the *χ* parameter, referred to as the aperiodic component, is computed as the negative slope of the log-log power spectrum (1/f slope). The aperiodic component is additionally parameterized with an ‘offset’ parameter, which reflects a shift of power spectrum. Since power spectrum normalization modifies relative differences in the offset of power spectrum, we did not include this parameter in our study to avoid misinterpretation of results.

### 2.3. Cortical network model and computation of field potentials

The cortical network model included an excitatory and an inhibitory population of leaky integrate-and-fire (LIF) spiking neuron models that interact through recurrent connections (P. Martínez-Cañada et al. 2021; Trakoshis et al. 2020; Mazzoni et al. 2015; Brunel and Wang 2003) (Fig. 1 A). Each population received two different Poisson signals: an external input with constant-rate (*ν*_0_) that can represent sensory or cortico-cortical inputs, and a noise input that captures intracortical fluctuations of neural activity. We used the same network configuration with the same parameters as described in previous publications (P. Martínez-Cañada et al. 2021; Trakoshis et al. 2020; Cavallari, Panzeri, and Mazzoni 2014). Briefly, the network consists of 5000 conductance-based LIF neuron models, 80% of which are excitatory (i.e., they form AMPA-like excitatory synapses with other neurons) and 20% are inhibitory (forming GABA-like synapses). Neurons in the network are randomly connected with each other (connection probability = 0.2). To account for neural activity with different E/I ratios, we performed several simulations of the recurrent network model where we systematically varied the ratio between excitatory and inhibitory synaptic conductances (*g_E_/g_I_*) across the range 0.06 – 0.2 (Fig. 1 B). We additionally set *ν*_0_ = 3 spikes/s, a low input strength that drives the network into an spontaneous activity regime. We repeated simulations for a given value of *g_E_/g_I_* with different random initial conditions (e.g., recurrent connections of the network). All simulations were performed using NEST v2.16.0 (Linssen 2018) and parallelized in a high-performance computing server (32-core CPU and 128 GB RAM).

**Fig. 1.**
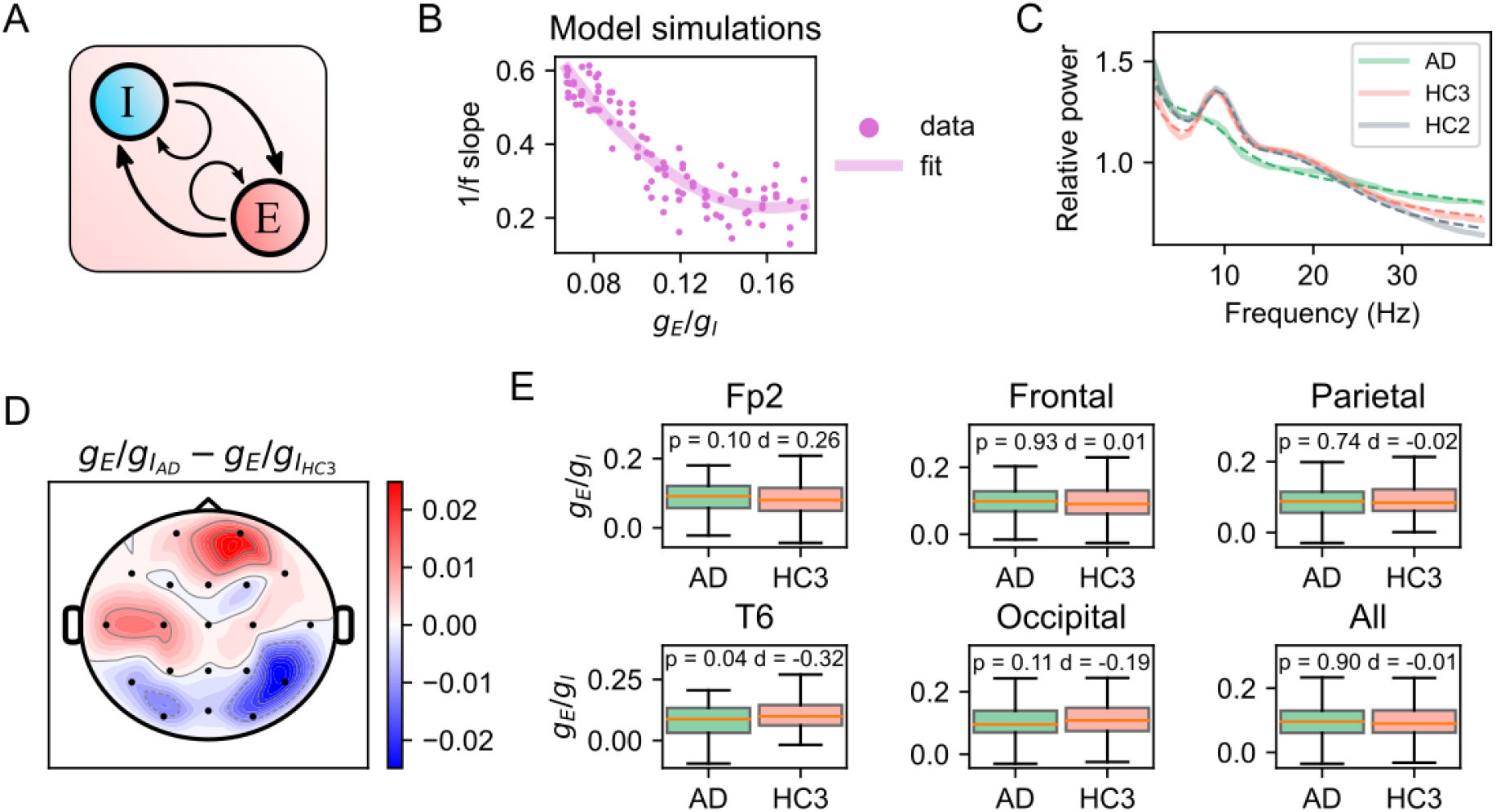
Changes of spectral properties in AD and E/I imbalance predictions from the recurrent network model. (A) Overview of the network model that includes recurrent connections between two types of populations: excitatory cells (E) and inhibitory cells (I). (B) 1/f slopes from model simulations as a function of the ratio between synaptic conductances (*g_E_/g_I_*) and fit of the polynomial regression model. (C) Normalized power spectrum across all individuals from the HC2, HC3 and AD groups and over all electrode locations. Dash lines correspond to spectral fits including aperiodic and periodic components. (D) Topographic representation of mean differences in *g_E_/g_I_*. (E) Predictions of changes in *g_E_/g_I_* for specific subsets of electrodes (p and d indicate p-value of the two-sided t-test and Cohen’s effect size respectively).

Field potentials cannot be directly computed from LIF neuron models because in a LIF model all transmembrane currents collapse into a single point in space and the resulting extracellular potential (and in turn, the resulting field potential) is zero. An approximation of field potentials can be calculated using variables available from simulation of the network model (e.g., synaptic currents). Here we computed an approximation of M/EEG signals as the sum of absolute values of AMPA and GABA postsynaptic currents on excitatory cells (∑|*I_AMPA_*| + ∑|*I_GABA_*|). This simple approach was shown to perform remarkably well in predicting simulated field potentials at different spatial scales, from local field potentials (LFPs) (Mazzoni et al. 2015) to EEGs (P. Martínez-Cañada et al. 2021) and in capturing more than 90 % of variance of empirical data recorded in neocortex (Barbieri et al. 2014; Mazzoni et al. 2008).

### 2.4. Regression model for estimating E/I ratio and statistical tests

We computed a least-squares polynomial regression model (degree = 2) with data from the whole set of simulations of the recurrent neural network (varying *g_E_/g_I_* across the range 0.06 – 0.2 and *ν*_0_ = 3 spikes/s). The regression model was computed using x = 1/f slopes from simulated field potentials and y = *g_E_/g_I_* values. Estimated parameters of the regression model were used to predict changes in *g_E_/g_I_* from slopes of empirical M/EEG data. Measures of slopes and predictions of *g_E_/g_I_* followed a normal distribution and had equal variances. We determined whether there was a significant difference between two groups of measures (e.g., control vs. AD) using a two-sided t test. Significant effects between more than two groups of measures (e.g., control, MCI converted and non-converted) were tested using a one-way ANOVA test and post-hoc pairwise comparisons with Tukey HSD confidence intervals (alpha = 0.05). We used Cohen’s d to account for effect size. In all statistical tests, data were pooled over all epochs and individuals.

## 3. Results

### 3.1. Opposing E/I shifts in posterior and anterior cortical areas of patients with AD

We first investigated whether variations in spectral slopes could be related to changes in E/I using computational modelling. We performed several simulations of a LIF neural network model with interacting excitatory and inhibitory neuronal populations (P. Martínez-Cañada et al. 2021; Trakoshis et al. 2020; Mazzoni et al. 2015) (Fig. 1 A) in which we systematically varied the ratio between synaptic conductances (*g_E_/g_I_*) (Fig. 1 B). From this model, we computed the network’s M/EEGs as the sum of absolute values of all synaptic currents, an approximation that has demonstrated to capture the main properties of field potentials at different spatial scales and has been validated on both simulated and real cortical data (P. Martínez-Cañada et al. 2021; Mazzoni et al. 2015; Barbieri et al. 2014; Mazzoni et al. 2008). Consistent with previous work (Trakoshis et al. 2020; Gao, Peterson, and Voytek 2017), 1/f slopes computed from simulated field potentials decreased (flatter slopes) with increasing *g_E_/g_I_*. Using simulation data, we computed a polynomial regression model to fit the relationship between *g_E_/g_I_* and slope values (*R*^2^ = 0.77, Fig. 1 B). These modelling results indicate that changes in synaptic E/I ratio are reflected by and thus could be potentially inferred from the readout of spectral slope.

We next analysed EEG data from an open EEG dataset (Meghdadi et al. 2021) that included resting-state EEG recordings of healthy individuals across different ages (HC2 and HC3 groups) and patients with AD. Comparing power spectra of AD patients and age-matched controls (HC3), we found that spectral recordings of AD patients displayed, as described in various studies (for review, see (Maestú et al. 2021)), slower brain oscillatory activity with a prominent reduction of alpha (8-12 Hz) and beta (12-30 Hz) rhythms and increased delta/theta (0.5 – 8 Hz) band activity (Fig. 1 C). Additionally, we observed a flattening of power spectrum in HC3 (older) individuals (Fig. 1 C), as well as smaller slopes (Supp. Fig. 1 A), compared to HC2 (younger) group, which was consistent with the hypothesis of increasing neural noise in healthy aging (Tran et al. 2020; Voytek et al. 2015).

To test whether slopes can be used to predict changes in E/I, we computed 1/f slopes of individuals in AD and HC3 groups and used the regression model to produce *g_E_/g_I_* predictions from these empirical slope measures. The topographic representation of changes in *g_E_/g_I_* (Fig. 1 D) revealed an interesting pattern in AD with opposing effects suggesting a coexistence of excitation- and inhibition-dominated regions. This is consistent with Aβ and tau effects at advanced stages of the disease, in which Aβ and tau pathologies are coestablished and coexist (Harris et al. 2020). Importantly, we compared local differences in *g_E_/g_I_* across subsets of electrodes (Fig. 1 E) and found that some regions, such as prefrontal locations (Fp2), exhibited strong shifts in *g_E_/g_I_* toward excitation (p = 0.1, d = 0.26), while *g_E_/g_I_* in temporal electrode locations (T6) showed significant shifts toward inhibition (p = 0.04, d = −.32). Differences in *g_E_/g_I_* between AD patients and controls were much weaker and insignificant when pooling data from all electrode locations (‘All’, Fig. 1 E), which suggests that local synaptic excitatory and inhibitory drives cancelled each other out at the global network level. Direction and strength of changes in slopes (Supp. Fig. 1 B and C) were largely in agreement with the patterns of differences in *g_E_/g_I_* but less prominent at Fp2 and T6 electrode locations.

**Supp. Fig. 1, related to Fig. 1.**
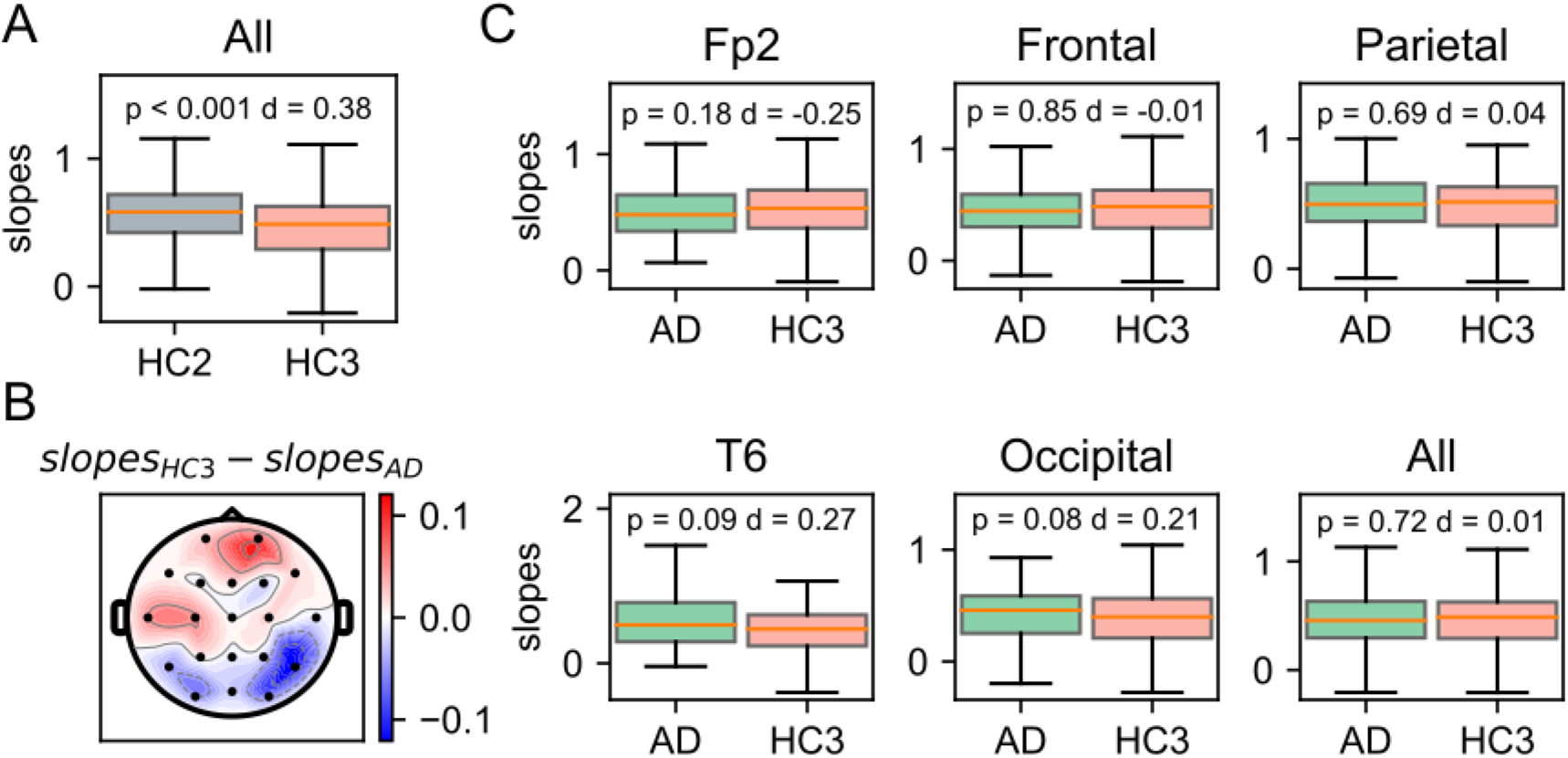
Shifts in 1/f slopes in aging and AD. (A) 1/f slope values by age pooled across all electrode locations and individuals. (B) Topographic representation of mean differences in slopes. (C) Comparison of slopes between AD and control (HC3) groups, computed using the same subsets of electrodes as shown in Fig. 1 E. In all statistical tests, p and d indicate p-value of the two-sided t-test and Cohen’s effect size respectively.

### 3.3. Cortex-wide hyperexcitability in MCI patients

We examined MEG data from a different publicly available dataset, the BioFIND dataset (Vaghari et al. 2022). This dataset included resting-state MEG data of MCI patients and healthy controls. Additional labels were provided for MCI patients indicating whether they subsequently progressed to probable AD (converted) or did not convert to a dementia diagnosis (not converted). We computed average MEG power spectra and found substantial differences in spectral properties between controls (‘Ctrl’) and MCI patients (Fig. 2 A). Power spectrum shapes of controls and not-converted MCI patients (‘MCI_n_’) largely overlapped across a broadband of low frequencies from 0.5 to 12 Hz. In contrast, the power spectrum of converted MCI patients (‘MCI_cv_’) comparatively captured some of the defining features of AD at lower frequencies, such as attenuation of alpha rhythm and increased delta/theta power. At higher frequencies (> 25 Hz), power spectra of MCI_n_ and MCI_cv_ eventually converged and showed higher power than control spectra.

**Fig. 2.**
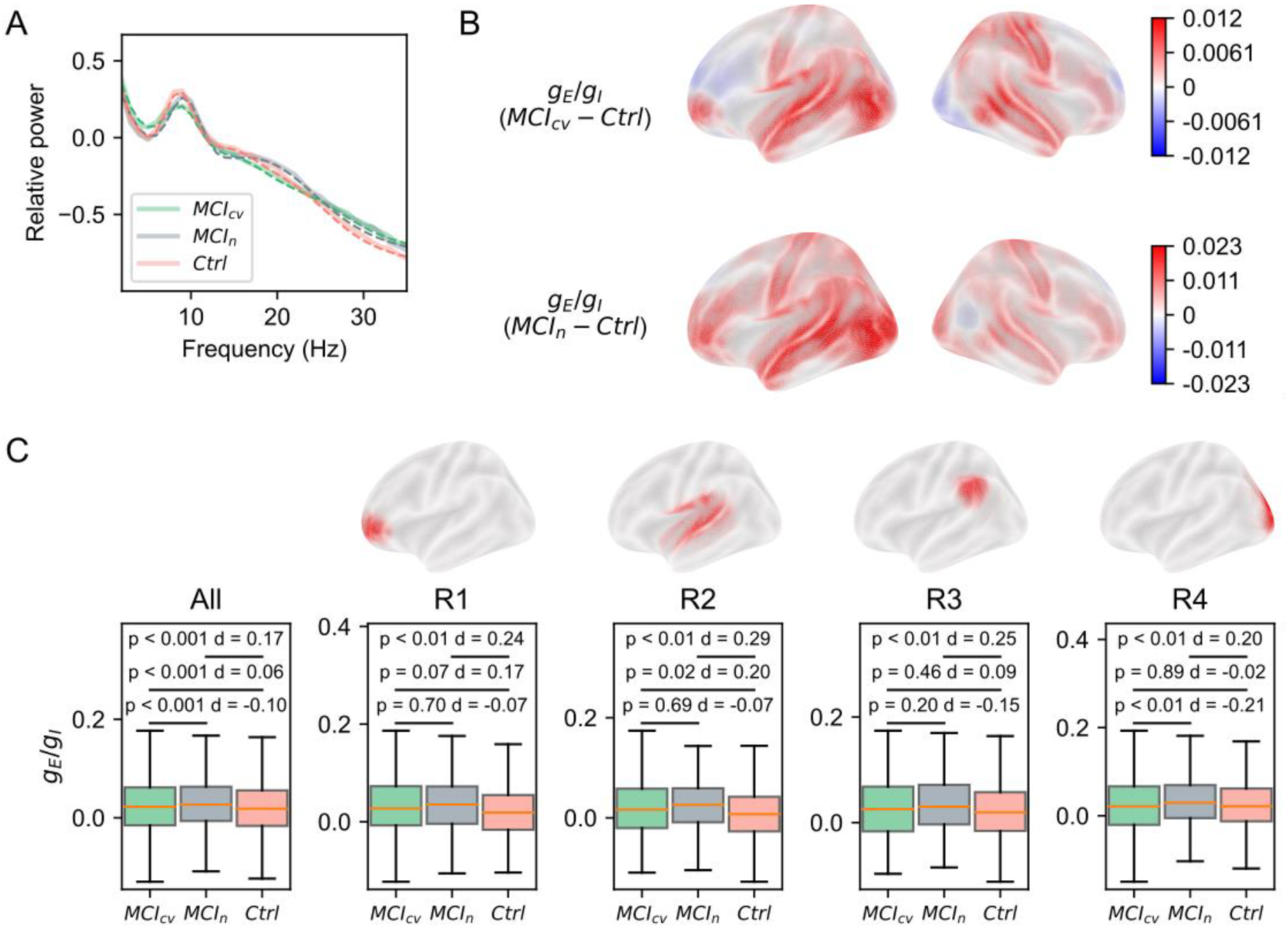
Spectral changes and E/I shifts in MCI. (A) Normalized power spectrum across all individuals from the control (Ctrl), MCI_n_ and MCI_cv_ groups and over all ROIs. (B) Surface plots (lateral view) representing mean differences in *g_E_/g_I_*. (C) Predictions of *g_E_/g_I_* pooled across all ROIs (‘All’), and for a subset of selected ROIs (R1, R2, R3 and R4) represented in the upper surface plots (p and d indicate p-value of the post-hoc Tukey HSD test and Cohen’s effect size respectively).

We then computed 1/f slopes (Supp. Fig. 3) and their corresponding *g_E_/g_I_* estimates (Fig. 2 B and C, and Supp. Fig. 2) for every one of the 38 ROIs of the parcellation used here (described in ref. (Colclough et al. 2015)). Hyperexcitability (i.e., increased *g_E_/g_I_*) ostensibly dominated along cortical regions in both MCI groups and agreed with clinical data of Aβ hyperactivity in early AD (Harris et al. 2020). Consistently, slopes of MCI_n_ and MCI_cv_ were overall reduced across ROIs (Supp. Fig. 3). The analysis of *g_E_/g_I_* predictions pooled across all ROIs (‘All’, Fig. 2 C) revealed statistically significant global differences between controls and MCI_n_, and between controls and MCI_cv_ (p < 0.001 in both cases) although small size effects (d ≤ 0.17). We finally compared regional *g_E_/g_I_* variations across subsets of ROIs (R1, R2, R3 and R4, Fig. 2 C) and observed that most *g_E_/g_I_* shifts of MCI_cv_ with respect to controls were insignificant or marginally significant, while *g_E_/g_I_* variations between MCI_n_ and controls were overall stronger and significant (p < 0.01, d ≥ 0.2 in all subsets of ROIs). The weaker increase in local and global excitability of MCI converters may be linked to the presence of inhibitory influences that may reflect a more advanced stage of the disease and the onset of tau pathology.

**Supp. Fig. 2, related to Fig. 2.**
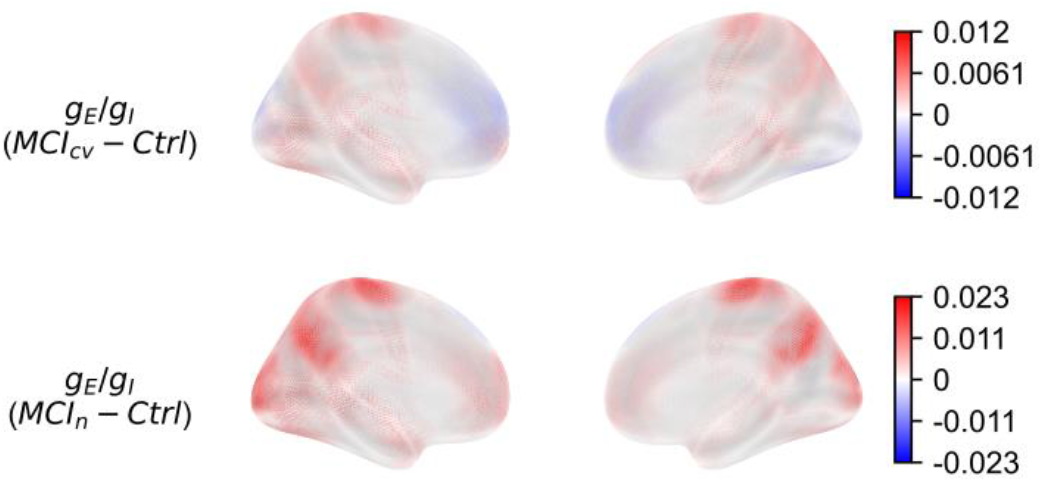
Surface plots (medial view) representing mean differences in *g_E_/g_I_*.

**Supp. Fig. 3, related to Fig. 2.**
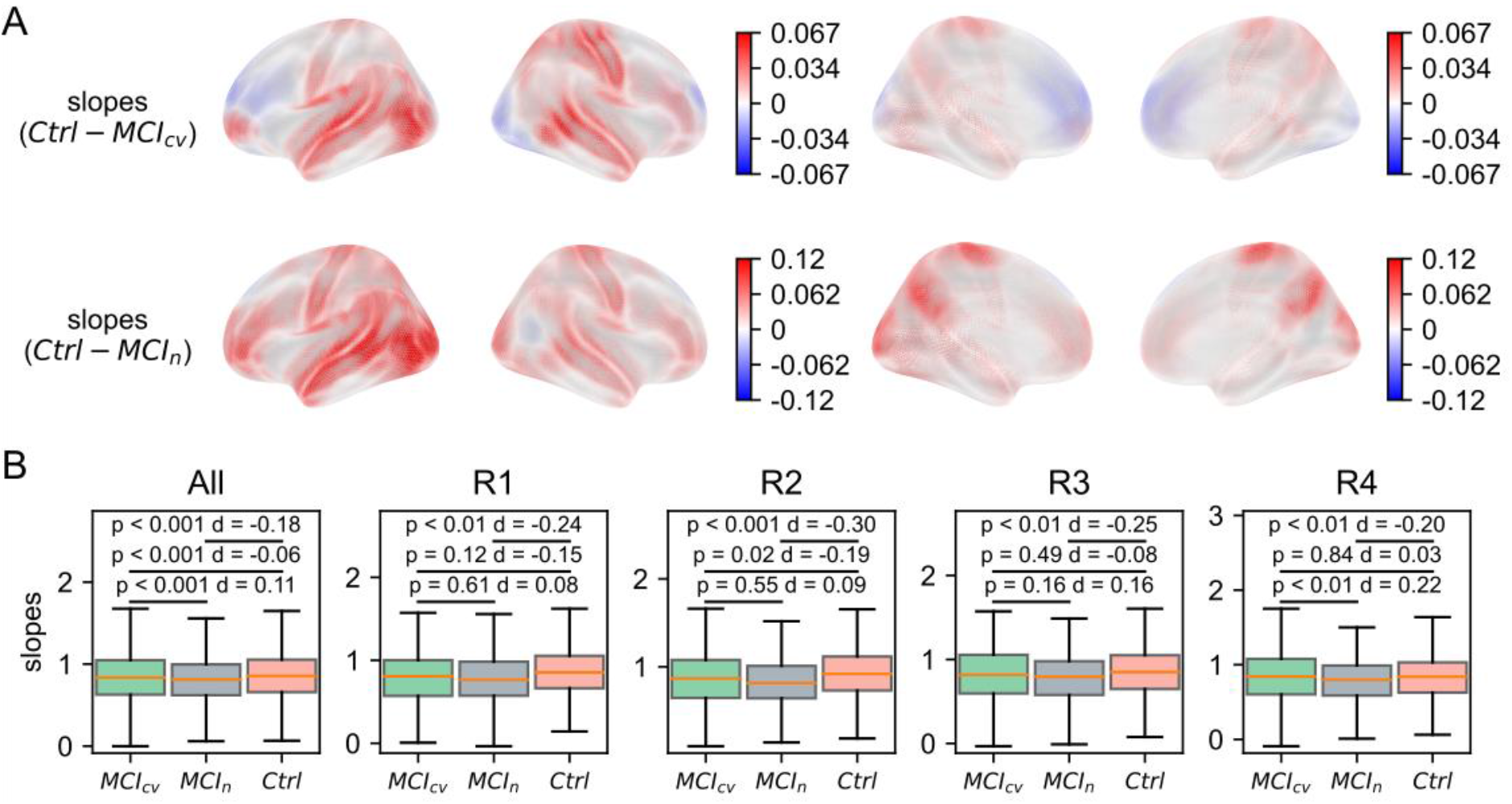
Changes of 1/f slopes in MCI. (A) Surface plots (lateral and medial views) of mean differences in slopes. (B) Comparison of slopes across the same subsets of ROIs as shown in Fig. 2 C.

## 4. Discussion

We have described a new method for inferring alteration in the E/I balance of neuronal circuits in Alzheimer’s disease using EEG and MEG measures. It is based on a recent neuroimaging biomarker, the 1/f slope, which has been proposed as a robust proxy marker for noninvasive real-time interrogation of E/I imbalance in many neuropsychiatric conditions (Ahmad et al. 2022). We used a regression model to map empirical 1/f slopes onto E/I estimations obtained from computational simulations of a recurrent cortical microcircuit model of spiking neurons that has shown to capture the main properties of simulated and real field potentials (P. Martínez-Cañada et al. 2021; Trakoshis et al. 2020; Mazzoni et al. 2015). We evaluated the ability of our approach to identify potential excitation-inhibition imbalances using recordings of patients at prodromal and clinical stages from publicly available EEG (Meghdadi et al. 2021) and MEG (Vaghari et al. 2022) datasets. Collectively, our results support the hypothesis that E/I imbalance in AD is reflected by and thus could be inferred from the readout of spectral slope and demonstrate the potential of this biomarker to be used to investigate circuit dysfunction in Alzheimer’s disease models. We discuss the implications and limitations of our method below.

### 4.1. Predictions of E/I imbalances are potentially associated with spatiotemporal progression of tau and amyloid-beta in AD pathophysiology at circuit-level

Aβ and tau dynamically modulate neuronal and circuit activity during AD progression (Maestú et al. 2021; Harris et al. 2020; Busche and Hyman 2020; Lewandowski, Maldonado Weng, and LaDu 2020; van der Kant, Goldstein, and Ossenkoppele 2020). At earlier stages, Aβ plaques accumulate and predispose neuronal circuits toward neuronal hyperexcitability (Fig. 3). Importantly, epileptic activity and enhanced seizure susceptibility are phenomena often observed in early AD (Cretin et al. 2016; Vossel et al. 2013). A key finding from the current study is consistent with these observations. We found that E/I prediction results from the cohort of patients with MCI, a prodromal state of AD (likely between early and mid stages in Fig. 3), consistently displayed a generalized pattern of hyperexcitability along cortical regions. Tau pathology, which is primarily associated with suppression of neuronal activity, by contrast, becomes evident in later stages of AD progression, at advancing disease stages (van der Kant, Goldstein, and Ossenkoppele 2020). Indeed, it has been shown that increase in tau pathology correlates much more strongly than Aβ pathology with neurodegeneration and cognitive impairment, the hallmarks of the clinical stage of AD. Tau aggregates are commonly found in the medial temporal lobe, starting in the parahippocampal gyrus, which includes the entorhinal cortex, from which they spread to limbic areas, and from there to the association areas (Busche and Hyman 2020). At advanced stages of the disease, thus, Aβ and tau pathologies coexist and interact, and their effects, in combination, can shift neuronal circuits from balanced E/I toward hyperexcitability or hypoexcitability. Our findings based on E/I estimations using EEG data from patients with AD draw remarkable potential parallels to the spatiotemporal evolution profiles of Aβ and tau between probably occurring at mid and late stages (Fig. 3). First, topographic representation of E/I estimates revealed a coexistence of excitation- and inhibition-dominated regions. Second, we found significant patterns of hyperexcitability in prefrontal locations (regions associated with initial Aβ deposition) and of hypoexcitability in the temporal lobe (associated with initial tau accumulation). Taken together, these results show the potential of our inference method to track E/I disruptions during AD progression both spatially and over time.

**Fig. 3.**
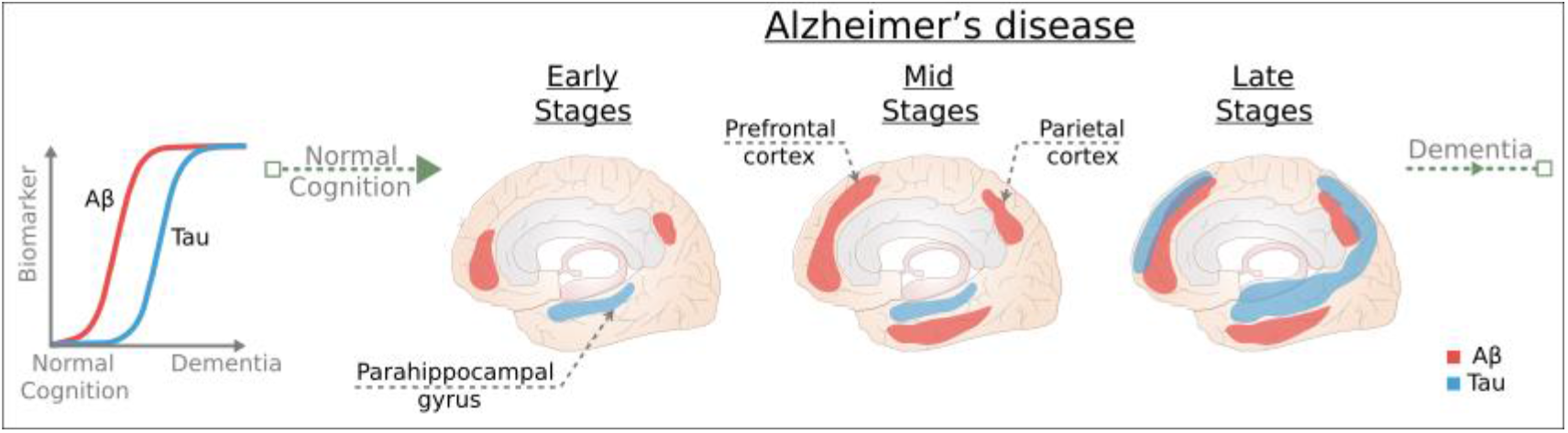
Schematic representation of the spatiotemporal progression of Aβ and tau biomarker levels during AD progression. As the disease progresses, Aβ and tau show different temporal evolution profiles in which Aβ plaque deposition precedes cortical tau pathology. Aβ and tau pathologies initially start in different brain regions. Aβ plaques are initially deposited in medial prefrontal and medial parietal regions, while tau aggregates first become evident in the medial temporal lobe, probably starting in the entorhinal cortex. During AD progression, Aβ and tau spread through different areas of cortex and may interact synergistically. Blue and red shaded areas indicate the areas affected by tau and Aβ pathology, respectively. This schematic plot was created based on refs. (Harris et al. 2020; Busche and Hyman 2020; van der Kant, Goldstein, and Ossenkoppele 2020; Leuzy et al. 2019).

### 4.2. Impact of our method to investigate brain dysfunction at the circuit level and monitor responses to drug treatments that aim to restore the balance between excitation and inhibition

A leading theory holds that many neurodevelopmental disorders, from autism to schizophrenia, and neurodegenerative diseases, such as Alzheimer’s disease, are the result of an imbalance in the activity of excitatory and inhibitory neurons (Maestú et al. 2021; Harris et al. 2020; Sohal and Rubenstein 2019; Lee, Lee, and Kim 2017; Nelson and Valakh 2015; Gogolla et al. 2009). Brain dysfunction has been traditionally investigated and diagnosed entirely at the behavioural level, whereas interventions aimed to correct alterations pharmacologically focused on the molecular level. A recent viewpoint argues that the most effective scale for neuroscientific investigation and intervention could be the circuit level (Maestú et al. 2021; Harris et al. 2020; Sohal and Rubenstein 2019; Palop and Mucke 2016), which is the intermediary scale bringing the gap between molecular and genetic changes at the microscopic level and macroscopic alterations in behaviour. Our modelling approach provides a mechanistic framework linking abnormal E/I ratio at the level of circuits (where neuronal and synaptic changes mediated by AD-related proteins possibly converge) to properties of macroscopic brain signals (M/EEG). Results of this study indicate that computational modelling at the circuit level could help integrate neurophysiological aspects of AD at the micro-, meso- and macroscale. Moreover, we have shown that our inference method largely tracks directionality of expected E/I shifts during AD progression and, thus, our approach could be used to estimate changes in excitatory and inhibitory circuit parameters in response to administration of pharmacological drugs commonly used for treatment of different clinical disorders (e.g., memantine, ketamine or lorazepam).

### 4.3. Novel noninvasive E/I proxy markers

Recently, a range of novel E/I biomarkers has been described that are noninvasive and applicable in humans and which can be deployed on a large scale. (Ahmad et al. 2022). These markers represent a key translational link across species and recording scales and can be applied in resting-state experiments and tasks that do not require extensive cognitive processing. They have been used to make inferences of E/I disruptions across many psychiatric and neurodevelopmental disorders, often combined with pharmacological or chemogenetic manipulations that modulate the level of inhibition and excitation (Manyukhina et al. 2022; Trakoshis et al. 2020; Molina et al. 2020), or to uncover the neural circuit basis of some key brain processes such as development (Chini, Pfeffer, and Hanganu-Opatz 2022; Schaworonkow and Voytek 2021) or aging (Tran et al. 2020; Voytek et al. 2015). Some of these novel E/I markers are aperiodic 1/f signal (Trakoshis et al. 2020; Gao, Peterson, and Voytek 2017), neuronal avalanches (Varley et al. 2020; Lombardi, Herrmann, and de Arcangelis 2017), long-range temporal correlations (LRTC) (Bruining et al. 2020), neural entropy (Yang et al. 2012; Shew et al. 2011) and microstates (Fingelkurts et al. 2004). Although each of these markers has been mapped to a different level of neural inference in the brain, they are likely intercorrelated with one another, and potentially estimate overlapping aspects of the same underlying signal at different scales. Further work would be needed to examine the relationship between neural variables measured by each biomarker and to investigate whether they can be used in combination for a more comprehensive interrogation of E/I balance. Additionally, combining these biomarkers with other imaging techniques, such as MRS and PET (Ranasinghe et al. 2022; Vaghari, Kabir, and Henson 2022; Popescu et al. 2020), will allow us to gain a better insight into the mechanisms of E/I by leveraging the strengths of each method.

### 4.4. Biophysical modelling in AD research

Previous models in AD research have lumped neural populations at each brain region into neural masses (neural mass models) (Ranasinghe et al. 2022; Monteverdi et al. 2022; van Nifterick et al. 2022; Stefanovski et al. 2021; Stefanovski et al. 2019; Zimmermann et al. 2018; de Haan et al. 2017). Using neural mass models, recent studies have found a relationship between abnormal excitatory and inhibitory time-constants and spatial depositions of Aβ and tau (Ranasinghe et al. 2022; Stefanovski et al. 2019), have linked neuronal hyperactivity in preclinical AD to oscillatory slowing (van Nifterick et al. 2022; Stefanovski et al. 2019) and have developed a successful strategy to preserve network integrity during AD progression (de Haan et al. 2017). A recent work has also used neural mass models to extract information about the excitatory/inhibitory balance in single subjects and suggested that AD subjects were characterized by increased global coupling and increased inhibition (Monteverdi et al. 2022).

Neural mass models have been proven useful to explain large-scale neuronal processes and structure-function coupling at macroscopic scales. Spiking and multicompartment models rather focus on describing neural phenomena at the micro- and mesoscopic scales and allow researchers to simulate synaptic connectivity, individual spike events and/or whole-cell dynamics, as well as heterogeneous parameter distributions in network populations. Due to limited computational resources and the higher level of complexity of point neuron models and especially multicompartment models, appropriateness and interpretability of these models at whole-brain scale has been questioned. However, these models are uniquely positioned at an intermediary scale of biophysical description, which facilitates interpretation of the relationship between neuronal and synaptic changes mediated by molecular and cellular mechanisms with circuit-level dynamics and properties of field potentials. Moreover, a range of studies have developed biophysics-based methods for computing ‘proxies’ of field potentials that provide accurate approximations of extracellular signals from simulations of spiking network models (Hagen et al. 2022; P. Martínez-Cañada et al. 2021; Hagen et al. 2018; Mazzoni et al. 2015). Recently, large-scale network models of cortex incorporating point neuron or multicompartment models and data-driven long-range connectivity between areas have shown to reproduce both experimental spiking statistics and cortico-cortical interaction patterns measured in functional MRI (Billeh et al. 2020; Schmidt et al. 2018). Spiking and multicompartment models offer a new perspective for unifying local and large-scale accounts of cortical dynamics and provide a promising platform for future studies of brain function and dysfunction.

### 4.5. Limitations

Our findings should be considered in the context of the following limitations. Our approach is based on a spiking network model of recurrently connected excitatory and inhibitory neuronal populations that has been constructed to approximate circuit phenomena with high accuracy at the local scale level (Cavallari, Panzeri, and Mazzoni 2014; Mazzoni et al. 2011; Mazzoni et al. 2010; Mazzoni et al. 2008). However, we did not model brain structure and function at the scale of macroscopic networks, that is, among brain regions and their long-range cortico-cortical connections. Mean field or neural mass models (Sanz Leon et al. 2013; Deco, Senden, and Jirsa 2012; Destexhe and Sejnowski 2009; El Boustani and Destexhe 2009), coupled using anatomic connectivity information, have been employed to account for whole-brain macroscopic phenomena, such as the emergence of patterns of coherent activity across regions in the cerebral cortex and propagation of slower oscillatory rhythms. The most effective scale for studying AD (and other clinical conditions) and for intervention remains an area of active research (Maestú et al. 2021; Harris et al. 2020). The challenge for future research on neural functional impairment will be to devise and implement the necessary multilevel computational modelling approaches that can integrate local circuitry and long-range circuit dynamics (Billeh et al. 2020; Schmidt et al. 2018; X.J. Wang 1999).

Another limitation of our approach is that the cortical circuit model in its current form does not include some important synaptic mechanisms such as NMDA synapses. NMDA receptors are critically involved in persistent cortical activity underlying working memory and evidence accumulation (Murray et al. 2014; X.J. Wang 1999). Disruptions in cortical NMDA receptor signalling have been associated with certain brain disorders, including AD (R. Wang and Reddy 2017), and likely affect E/I balance (Molina et al. 2020; Foss-Feig et al. 2017; Driesen et al. 2013; Fellous and Sejnowski 2003). Using our simplified model of recurrent AMPA and GABA synaptic currents, we have demonstrated here that E/I imbalances across cortical regions can be consistently associated to spatiotemporal evolution profiles of AD-related proteins. An interesting topic for a future study would be to extend the spiking network model to include NMDA synapses and to assess their impact on the current prediction results of E/I alterations in AD.

Finally, it is noteworthy to mention that the experimental protocols for recording the open EEG and MEG data used in this study and for recruiting participants, though they are out of our control, include additional limitations. First is the fact that the data come from different sites. Although the recording machine models were identical, there are still differences in individual devices (e.g., in their noise level). The clinical assessments followed international guidelines but the precise categorization as AD, MCI and control is likely to differ across clinicians at different sites. The tasks performed prior to resting were different and likely had different levels of fatigue, which could have contributed to noise in the data. The groups also differ in other respects, such as age and education.

## 5. Conclusion

A prominent view is that circuit-level effects mediated by AD-related proteins provide diagnostic and prognostic information and offer a unique framework to understand AD and develop effective therapeutic interventions. Disambiguating the complexity of Aβ and tau excitatory and inhibitory influences and their interacting effects at the synaptic and circuit scales will be a crucial next step to identify rational therapeutic targets in AD. The approach presented here, consisting of recurrent network modelling and multimodal neuroimaging data, provides a general framework for investigating circuit-level excitatory/inhibitory imbalances and could help in linking micro- and macroscopic scales of investigation in AD.

## Code and data availability

Code used to compute 1/f slopes and E/I predictions, as well as code supporting the findings of this study can be found at https://github.com/pablomc88/EEG-and-MEG-Proxy-Marker-of-Excitation-Inhibition-Imbalance-in-Alzheimer-s-Disease. The EEG dataset was downloaded from Mendeley Data and is accessible via the following DOI: 10.17632/wttpypkctg.2. The MEG dataset used in this study is available on request from the Dementia Platform UK (DPUK) (https://portal.dementiasplatform.uk/Apply).

## Acknowledgements

We thank members of the NeuroEngineering and Computing (NECo) Lab (University of Granada, Spain) for their invaluable discussions and feedback on this work.

This work was conducted using the MRC Dementias Platform UK (DPUK). DPUK is a Public Private Partnership funded by the Medical Research Council (MR/L023784/1 and MR/009076/1). For further information on this resource visit www.dementiasplatform.uk.

## Author contributions

**Pablo Martínez-Cañada**: Conceptualization, Methodology, Formal analysis, Software, Writing - original draft, Writing - review & editing, Funding acquisition; **Eduardo Perez-Valero**: Methodology, Writing - review & editing, Funding acquisition; **Jesus Minguillon**: Methodology, Writing - review & editing, Funding acquisition; **Francisco Pelayo**: Methodology, Writing - review & editing, Funding acquisition; **Miguel A. López-Gordo**: Methodology, Writing - review & editing, Resources, Funding acquisition; **Christian Morillas**: Methodology, Writing - review & editing, Resources, Funding acquisition.

